# Modeling Molecular Pathogenesis of Idiopathic Pulmonary Fibrosis-Associated Lung Cancer in Mice

**DOI:** 10.1101/2023.06.20.545616

**Authors:** Ivana Barravecchia, Jennifer M. Lee, Jason Manassa, Brian Magnuson, Sophia Cavanaugh, Nina G. Steele, Carlos Espinoza, Craig J. Galban, Nithya Ramnath, Timothy L. Frankel, Marina Pasca di Magliano, Stefanie Galban

**Author notes:** **Corresponding Author:** Stefanie Galban Department of Radiology, Center for Molecular Imaging University of Michigan, 109 Zina Pitcher Place, Ann Arbor MI 48109-2200 USA. Co-first author (Ivana Barravecchia, Jennifer M. Lee, Jason Manassa). **Financial Support:** This work was supported by the University of Michigan Rogel Cancer Center “First and Goal” funding to the Galban Lab. Flow cytometry work was performed at the University of Michigan Flow Cytometry Core, which is supported by the National Cancer Institute (NCI) of the National Institutes of Health (NIH) Grants P30 CA046592 and P30 CA04659229. Histology was performed at the Rogel Cancer Center Tissue and Molecular Pathology Shared Resource, which is funded by NCI Grant P30 CA04659229. Multiplex immunohistochemistry (mIHC) was performed by the Frankel Lab and supported by the National Institute of Diabetes and Digestive and Kidney Diseases (NIDDK) of the NIH Grant R01 DK128102. CyTOF was performed at the Flow Cytometry Core at the University of Rochester Medical Center, with guidance and analysis provided by the Pasca di Magliano Lab, which is supported by NCI Grants R01 CA260752, R01 CA271510, U01 CA224145, U01 CA274154, and U54 CA274371. **Conflicts of Interest**: The authors declare no potential conflicts of interest.

## Abstract

Idiopathic Pulmonary Fibrosis (IPF) is characterized by progressive, often fatal loss of lung function due to overactive collagen production and tissue scarring. IPF patients have a sevenfold-increased risk of developing lung cancer. The COVID-19 pandemic has increased the number of patients with lung diseases, and infection can worsen prognoses for those with chronic lung diseases and disease-associated cancer. Understanding the molecular pathogenesis of IPF-associated lung cancer is imperative for identifying diagnostic biomarkers and targeted therapies that will facilitate prevention of IPF and progression to lung cancer. To understand how IPF-associated fibroblast activation, matrix remodeling, epithelial-mesenchymal transition, and immune modulation influences lung cancer predisposition, we developed a mouse model to recapitulate the molecular pathogenesis of pulmonary fibrosis-associated lung cancer using the bleomycin and the Lewis Lung Carcinoma models. Models of pulmonary fibrosis, particularly bleomycin-induced fibrosis, do not recapitulate all aspects of human disease; however, to simplify nomenclature, we refer to our bleomycin-induced fibrosis model as IPF. We demonstrate that development of pulmonary fibrosis-associated lung cancer is linked to increased recruitment or reprogramming of tumor-associated macrophages and a unique gene signature that supports an immune-suppressive microenvironment through secreted factors. Not surprisingly, pre-existing fibrosis provides a pre-metastatic niche and results in augmented tumor growth. Tumors associated with bleomycin-induced fibrosis are characterized by an epithelial-to-mesenchymal transition characterized by dramatic loss of cytokeratin expression.

**Implications:** We provide new therapeutic targets that may aid the characterization of tumors associated with lung diseases and development of treatment paradigms for lung cancer patients with pre-existing pulmonary diseases.

## Introduction

Idiopathic pulmonary fibrosis (IPF) is a debilitating and fatal lung disease with a median survival of 3-4 years following diagnosis [1, 2]. The approximately 3 million people worldwide affected by IPF exhibit signs of permanent lung scarring and suffer respiratory and other organ failure related to hypoxia [3]. Importantly, the global COVID-19 pandemic has intensified the seriousness of pulmonary fibrosis because lung infections, including COVID-19, worsen fibrosis in the lung [4]; furthermore, the effects of IPF on lung function increase patients’ risk of developing COVID-19, while those recovering from severe COVID-19 are at high risk for developing pulmonary fibrosis [5, 6]. While many genetic and environmental factors are linked to IPF, such as smoking and other forms of oxidative damage, the main cause of IPF remains largely unknown, thus preventing optimal treatment and diagnosis [7]. Treatment options for IPF remain limited and include anti-inflammatory therapy, transplant, palliation, or clinical trial recruitment [8]. However, recent advances in the field have led to discovery of novel anti-fibrotic agents like pirfenidone and nintedanib, which have been shown to significantly slow disease progression in IPF [9, 10]. These therapeutic options slow progression and temporarily alleviate the symptoms of fibrotic scarring. However, even with these treatments, IPF continues to progress, reflected by median survival rates of 3-4 years [1, 2].

While IPF is a devastating and often fatal inflammatory lung disease, lung cancer remains the leading cause of cancer related mortality worldwide [11]. The two main types of lung cancer, non-small cell lung cancer (NSCLC) and small cell lung cancer (SCLC), are characterized by different mutations, mutational loads, and survival rates. Around 80% of lung cancers are NSCLC [12], which are sub-grouped into adenocarcinomas, squamous cell carcinomas and large cell carcinomas. These subtypes differ at the molecular level, depending on whether mutations occur in *TP53, EGFR, KRAS, LKB1, PTEN,* or *BRAF* [13, 14]. Adenocarcinomas, which are often associated with *KRAS* mutations, are the most common form of lung cancer, accounting for 40% of all NSCLC [15]. The remaining 15-20% of lung cancers are SCLC; these are more aggressive, with a high mutational load and dismal survival rate, and mostly found in smokers [16]. With the emergence of immune checkpoint therapies and KRAS targeted therapies, great strides are being made in the treatment of lung cancer. However, an estimated 127,070 Americans will die of lung cancer in 2023, accounting for ∼21% of all cancer related deaths in the United States [17].

Epidemiological evidence indicates that IPF increases the risk of lung cancer by sevenfold [18] and thus, about 22% of patients with IPF develop lung cancer and exhibit a worse prognosis with poorer survival compared to the already dismal survival of IPF patients [19–22]. Cigarette smoke, chronic lung diseases like chronic obstructive pulmonary disease (COPD), fibrotic disorders, and recently, COVID, increase the risk for lung cancer in IPF patients [23, 24]. Since no guidelines exist to screen IPF patients for lung cancer, and common therapeutic options for lung cancer often exacerbate IPF, a critical need remains to identify the mechanistic link between these diseases, and in turn, illuminate the molecular underpinnings of IPF-associated lung cancer (IPF-LC) and the potential drivers that lead to lung cancer in 22% of IPF patients. To our knowledge, there is currently only one pre-clinical model that mimics the pathobiology of pulmonary fibrosis-associated lung cancer, described most recently in a study of tumor-associated fibrosis by Herzog and colleagues [25].

To gain a mechanistic understanding of the pathobiology of IPF-LC and identify and evaluate new therapeutic strategies, we developed a novel mouse model that combines the bleomycin-induced fibrosis model with the Lewis Lung Carcinoma model in a syngeneic background to recapitulate the human disease [26, 27]. Previous work has shown that mouse models of bleomycin-induced pulmonary fibrosis do not uniformly recapitulate human disease; in their study of three models, Gul and colleagues reported that a tail-vein injection model of bleomycin-induced fibrosis most closely recapitulates idiopathic pulmonary fibrosis [28]. Since our model is characterized by some aspects associated specifically with IPF, we discuss this disease throughout the paper. Because the tumor cells (LLC-1) are immunologically compatible with the host, the role of the immune response, which is a key contributor to cancer progression and metastasis, can be studied. This new murine model allowed us to interrogate the role of the tumor microenvironment (TME) in the context of fibrosis, lung cancer or pulmonary fibrosis-associated lung cancer and at the same time, provide mechanistic insight into the pathobiology of IPF and importantly, new therapeutic biomarkers and targets.

## Materials and Methods

### Animals

Female and male 8–10-week-old C57BL/6-albino mice (Jackson Laboratory, strain/stock: 000058 B6(Cg)-Tyr<c-2J>/J) were maintained in accordance with the University of Michigan’s Institutional Animal Care and Use Committee guidelines approved protocol (UCUCA PRO 00010349). C57B/L6-albino mice were randomized into 4 different experimental cohorts: (1) control mice not receiving bleomycin or LLC-1 Luciferase expressing cells (LLC-1 Luc), (2) mice that received two doses of 0.5mg/kg and 1 mg/kg bleomycin in 50 µl saline (double oropharyngeal aspiration (OA) schedule at day 0 and day 4) to induce fibrosis, (3) mice wherein LLC-1 Luc cells were injected intravenously through the tail vein (IV) at a concentration of 1×10^6 cells per mouse in 100 µl saline and (4) mice where LLC-1 Luc cells were implanted intravenously (IV) into bleomycin pre-conditioned mice (OA). Since the bleomycin model exhibits two distinct phases, the inflammatory (2 weeks) and the fibrotic (2 weeks) phase, LLC-1 Luc cells were injected 2 weeks post bleomycin administration during the fibrotic phase. We found that a double OA schedule of bleomycin at day 0 and day 4 produced the most consistent induction of lung fibrosis as previously described. [29, 30].

### Transgenic mouse model

Transgenic CCSP-rtTA mice [31] were first crossed with transgenic TRE-TGF-alpha mice [32]. The double transgenic offspring (CCSPrtTA+, TRE-TGFa+) of those was crossed to the transgenic TRE-KrasG12D model [33] to obtain four experimental groups: controls, IPF (CCSPrtTA+, TRE-TGFa+, TRE-KrasG12D-), LC (CCSPrtTA+, TRE-TGFa-, TRE-KrasG12D+) and IPF-LC (CCSPrtTA+, TRE-TGFa+, TRE-KrasG12D+). To induce IPF, LC or IPF-LC, mice were given doxycycline (Sigma, catalog number D9891) containing water at a concentration of 0.5 mg/ml at study start and when mice were 6 weeks and older. Two different ‘control groups’ were used for this study to control for the effects of doxycycline on lung tissue and for leakiness of the TetO inducible genes. Single transgenic control mice were given doxycycline in the drinking water at the same dose as experimental cohorts and triple transgenics were given regular drinking water.

### Bleomycin

Bleomycin was obtained through the University of Michigan Health Services Pharmacy services (Meitheal Pharmaceuticals, cat# 71288-106-10). Bleomycin powder was resuspended in 1xPBS at a concentration of 1 U/ml (equal to 1 mg/ml) and stored at −20°C. A dose of 0.5 mg/kg was given by oropharyngeal aspiration at the beginning of the study (day 0) and 1 mg/kg four days later (day 4).

### Cell lines

The murine Lewis Lung Carcinoma-1 (LLC-1) cell line was obtained from ATCC (ATCC® CRL-1642™) and maintained in DMEM with 10% FBS and 1% Pen/Strep. LLC-1 cells were infected with lentivirus FUGW [34] to express firefly luciferase and GFP to obtain LLC-1 Luciferase expressing cells for bioluminescence imaging (LLC-1 Luc). LLC-1 cells bear a heterozygous *Kras^G12C^* mutation [30].

### Bioluminescence Imaging

In vivo bioluminescence imaging (BLI) was performed under anesthesia using the IVIS Spectrum In Vivo Imaging System (PerkinElmer) according to manufacturer protocol. In short, mice received an intraperitoneal injection of 100 µl D-luciferin (stock of 40 mg/mL, Promega, E160). Mice were allowed to move freely for five minutes post injection prior to initiating anesthesia via isoflurane. Mice were imaged five minutes after isoflurane administration (10 minutes post injection). Imaging occurred at indicated timepoints post-implantation of LLC-1 Luc cells. Quantification of total flux (photons per second, p/s/cm^3^) using a standardized circular region of interest (ROI) spanning the entire lung was performed at each time point using IVIS Spectrum in vivo imaging software.

### Microcomputed tomography (CT imaging)

μCT imaging was performed at indicated timepoints using a Siemens Inveon System with the following parameters: 80 kilovolt peaks (kVp), 500 μA, 400-ms exposure, 360 projections over 360°, and 49.2-mm field of view (56-μm voxel size) as previously described [35]. Quantitative analysis was performed on automatically segmented lung volumes as the sum of lung, tumor, and vascular tissues. After image calibration to Hounsfield units (HU) using air and a water phantom, segmentation of the lungs was accomplished using a connected threshold algorithm developed in-house (MATLAB) with a threshold of −200 HU. This volume was subtracted from the total chest volume to approximate the tumor plus vascular volume. An assumption with this analysis is that vasculature should be similar between all subjects, with changes in this volume indicative of tumor volume changes [35].

### Multiplex immunohistochemistry and multispectral imaging

Multiplex immunohistochemistry or immunofluorescence (mIHC/IF) was performed as previously described [36]. In brief, five-micron murine lung sections were transferred onto charged slides and slides were baked at 60°C for 1 hour. This was followed by deparaffinization with xylene for 10 minutes in triplicate. Deparaffinization was followed by 100%, 90%, and then 70% ethanol application for 10 minutes each. Slides were washed in deionized water for 2 minutes followed by neutral buffered formalin for 30 minutes. The Opal 7 manual kit (PerkinElmer) was used according to manufacturer instructions. After each antigen retrieval, slides were stained with antigen-specific primary antibodies followed by Opal Polymer (secondary antibody). Application of the Opal TSA created a covalent bond between the fluorophore and the tissue at the site of horseradish peroxidase. Each antigen retrieval step was performed using either AR6 or AR9 antigen retrieval buffer, which allowed for the removal of prior primary and secondary antibody while the fluorophore remained covalently bonded to the tissue antigen. This allowed for use of the same host species antibody while also amplifying the signal. Antibodies used for this study can be found in **Supplemental Table 1.** Images of the stained murine lung sections were taken using the Mantra^TM^ Quantitative Pathology Workstation (PerkinElmer). One image per core was captured at 20x magnification. All cube filters were used for each image capture (DAPI, CY3, CY5, CY7, Texas Red, Qdot) and the saturation protection feature was utilized. After all images were acquired, images were analyzed using inForm® Cell Analysis^TM^ software (PerkinElmer). All images were batch analyzed and basic phenotypes were created using the in-Form training software. Basic phenotypes were composed of the following: Arg1, CK19, DAPI, CD3, F4/80, and CD8. The scoring feature was used to determine the appropriate range of the mean signal intensity of each stain within the cytoplasm for CD8 and nucleus for DAPI. These scoring ranges were used to make secondary phenotypes in R: macrophages, T-cells, epithelial cells, and ‘other’. Final phenotypes, including primary phenotypes (CK19, F4/80 and CD3) and secondary phenotypes made in R (Macrophages and CD8 positive T cells) were quantitatively analyzed using a program designed in R.

### Histology and immunohistochemistry

Murine lungs were perfused in 1x PBS then fixed in formalin and stored in 70% ethanol before embedding in paraffin. The top half of the right lung lobe was used for histology and sent to the University of Michigan Histology Core for paraffin embedding, sectioning, Hematoxylin and Eosin (H&E) and Masson Trichrome staining. Additional sections were hybridized with antibodies against Ki67, CD3, CD8 and CK7, performed by the core facility. In brief, for H&E staining, sections were deparaffinized in xylene, re-hydrated in ethanol, and briefly washed in distilled water. Staining was performed with Harris hematoxylin solution, and counterstained with eosin-phloxine solution, prior to mounting with xylene based mounting medium. For Masson Trichrome staining, Bouin’s solution was preheated for 1 hour at 56-60°C. Sections were deparaffinized and hydrated in deionized water. Sections were then incubated in Bouin’s solution for 1 hour in oven. Slides and lids were removed from the oven, allowed to cool for 10 minutes, then washed in tap water until their yellow color disappeared (approx. 10 minutes). Slides were placed in Weigerts Iron Hematoxylin solution for 10 minutes and rinsed in running tap water for 5 minutes. Rinse was repeated in deionized water. Subsequently, slides were blotted to remove excess water and stained in Biebrich Scarlet-Acid Fuchsin for 10 minutes, then rinsed in deionized water until clear. Slides were again blotted to remove excess water and incubated in phosphomolybdic-phosphotungstic acid for 15 minutes. Slides were blotted, incubated in aniline blue for 8 minutes and rinsed in deionized water until clear. Slides were dehydrated in 70% ethanol, 95% ethanol and 100% ethanol for 10 seconds each with agitation prior to mounting with xylene based mounting medium. Quantification of trichome staining was performed using Image J software.

### Evaluation of lung lesion size and number

H&E stained sections of the murine lungs were scanned using the Nikon Supercool Scan 5000 at 1X magnification. Evaluation of lung lesion number and size was blinded and performed on at least three sections per experimental group by three different readers. The number of lesions was counted, and the size measured with a ruler in cm on 1X magnified scans. The number of lesions and lesion size was graphed for each group and statistical significance was determined by one-way ANOVA using GraphPad Prism.

### Cytometry by Time-of-Flight (CyTOF)

Murine lung tissues from all four experimental cohorts were placed into DMEM Complete after lung perfusion with 1xPBS. Subsequently, lung tissues were mechanically minced and enzymatically digested with collagenase P (1 mg/mL DMEM) and subsequently filtered through a 40 µm mesh to obtain single cells. Up to 1 × 10^7^ cells were stained with Cell-ID cisplatin (1.67 μmol/L) for 5 minutes at room temperature, followed by a ‘Fix and Perm-Sensitive Surface Epitopes and Nuclear Antigen Staining’ protocol according to the manufacturer’s instructions (Fluidigm) for mouse samples [37]. Briefly, after quenching cisplatin reaction with 5X volume of MaxPar® cell staining buffer, cells were centrifuged at 300 × g for 5 minutes. Up to 3 million cells per sample were stained with cell surface antibody cocktail (see **Supplemental Table 2**) in 100 µl volume of MaxPar® cell staining buffer for 30 minutes at room temperature. Cells were subsequently washed twice in 1 ml MaxPar® cell staining buffer and then fixed in 1.6% freshly made formaldehyde solution for 10 minutes at room temperature. After fixation, cells were washed once in 1 ml MaxPar® cell staining buffer and permeabilized with 1 ml nuclear antigen staining buffer for 20 minutes at room temperature. Cells were then washed twice with 1 ml nuclear antigen staining perm and centrifuged in between at 800 × g for 5 minutes, followed by staining with intracellular antibody cocktail (see **Supplemental Table 2**) in 50µl volume of nuclear antigen staining perm for 45 minutes at room temperature. Subsequently, cells were washed with 2 ml nuclear antigen staining perm followed by a wash with 2 ml MaxPar® cell staining buffer, resuspended in 2 ml cell intercalation solution (125 nM Cell-ID Intercalator-Ir in Maxpar fix and perm buffer) and shipped to the Flow Cytometry Core at the University of Rochester Medical Center, where sample preparation was finalized, and Mass Cytometer analysis performed on a CyTOF2. Raw FCS files were analyzed using the Premium CytoBank Software (cytobank.org) and FlowSOM-viSNE [38] or viSNE as previously described [37]. Statistical analyses were performed in GraphPad Prism using paired t-tests with statistical significances established with a p-value less than 0.05.

### Flow Cytometry

Lung tissue was collected in 10 mL of FACS buffer (2% FBS, 500mL PBS, 5mM EDTA) and pushed through 70 µm cell strainer with syringe plunger. Red cell lysis was performed in 10 ml ACK (Ammonium-Chloride-Potassium) lysing buffer (Invitrogen, Cat # 11814389001) following manufacturer’s instructions. Cells were washed, centrifuged and 500,000 cells were stained in 200 µl 1x PBS plus 0.5 µg of indicated antibody for 30 min in the dark: APC anti-mouse CD274 (B7-H1, PD-L1, Biolegend #124311) or appropriate IgG isotype. After incubation 500 µl of the FACS buffer was added to the antibody staining solution and centrifuged for 1800 RPM for 5 minutes. Supernatant was removed and the process repeated for a total of two washes. After the second wash, the pellet was resuspended in 500 µl of FACS buffer and cells were analyzed using Accuri Flow Cytometer. Data was analyzed using Accuri software and data graphed using GraphPad Prism.

### Bulk RNA Sequencing

RNA was isolated from lung tissue using the Qiagen RNA isolation kit according to manufacturer protocol. RNA was stored in 30 µl of RNase/DNase free water and quantified using the NanoDrop 2000c (Thermofisher). 30 µg of RNA was sent to the University of Michigan Advanced Genomics Core for RNA Quant Sequencing. Total RNA quality was assessed using the Tapestation 4200 (Agilent). 500 ng of DNase-treated total RNA was used to generate QuantSeq 3’ mRNA FWD (Lexogen). Pooled libraries were subjected to 100 bp paired-end sequencing according to manufacturer protocol (Illumina NextSeq550). Bcl2fastq2 Conversion Software (Illumina) was used to generate de-multiplexed Fastq files. Reads were trimmed for adapters and quality using TrimGalore 0.5.0. Trimmed reads were aligned to the mm10 reference using STAR 2.6.0a. Annotations for mm10 were from UCSC and obtained from iGenomes (Illumina). Alignments were collapsed to UMIs using collapse_UMI_bam from Lexogen. Gene counting was carried out using Rsubread/featureCounts. DESeq2 1.38.3 [39] was used for differential gene expression analysis and genes with adjusted p-value < 0.05 and fold-change > 1.5 (log_2_ fold-change > 0.58) in either direction were considered significant. Genes uniquely regulated in the IPF-LC group were selected if they were significantly up- or downregulated in the IPF-LC group when compared to genes in the control group. The same genes had to show no statistically significant regulation in the LC or IPF group when compared to the control group. In other words, these genes were unique to the IPF-LC group when compared to normal tissue and to both IPF and lung cancer tissues. The datasets supporting the current study are available from the corresponding author upon request and available in the NIH Gene Expression Omnibus database (GEO). RNA raw data files and analysis was deposited with GEO accession number GSE224134.

## Results

### Pre-existing pulmonary fibrosis results in augmented tumor growth

Mouse models represent useful tools for advancing mechanistic understanding of diseases like lung cancer and testing new therapeutic strategies. To our knowledge, no murine model that recapitulates IPF-LC has been developed. In this newly developed IPF-LC mouse model, we induced lung fibrosis by intratracheal delivery of bleomycin and pulmonary lesions through intravenous injection of the murine, heterozygous *Kras^G12C^*mutant Lewis lung carcinoma cell line (LLC-1) in a syngeneic C57BL/6 background (Fig. 1A). Two distinct phases for bleomycin-induced IPF, an inflammatory (0-2 weeks) and a fibrotic (2-4 weeks) phase, have previously been described [29]. This guided our decision to inject LLC-1 Luc cells two weeks post instillation at the beginning of the fibrotic phase. We optimized the bleomycin oropharyngeal aspiration schedule and confirmed a previously reported benefit of double administration on day 0 and 4 to obtain the most consistent induction of lung fibrosis [40]. Development of pulmonary fibrosis and lesions were assessed by micro-CT and bioluminescence (Fig. 1B-1D), respectively, as well as by histological analysis at the end of the study (Fig. 1E). As expected, lung metastases were detected in the LC group (‘LC’), which received LLC-1 Luc cells alone, and in the IPF-LC group, where lung fibrosis was induced prior to LC initiation (‘IPF-LC’). Bioluminescence activity increased significantly over time in the IPF-LC group when compared to the LC group, indicative of more aggressive tumor seeding and growth in lungs due to pulmonary fibrosis (Fig. 1C). This finding was confirmed by histological analysis at the end of the study, when both the number and size of the pulmonary lesions were significantly augmented in the IPF-LC group versus the LC group (Fig. 1F and G). Statistical analysis between groups to assess the correlation between lung cancer and fibrosis revealed that mice with fibrosis experienced a higher tumor burden and larger lung tumors when compared to mice with no fibrosis (Fig. 1F and G). As expected, we also observed pulmonary fibrosis, determined by trichome staining of collagenous connective tissue fibers, in both bleomycin-induced fibrosis mice (‘IPF’) and the IPF-LC group (Fig. 1H). Interestingly, LLC-1 Luc cells were preferentially detected around the airways and the lung periphery, correlating with deposition of collagenous fibers post bleomycin administration in these areas (Fig. 1E).

**Figure 1:**
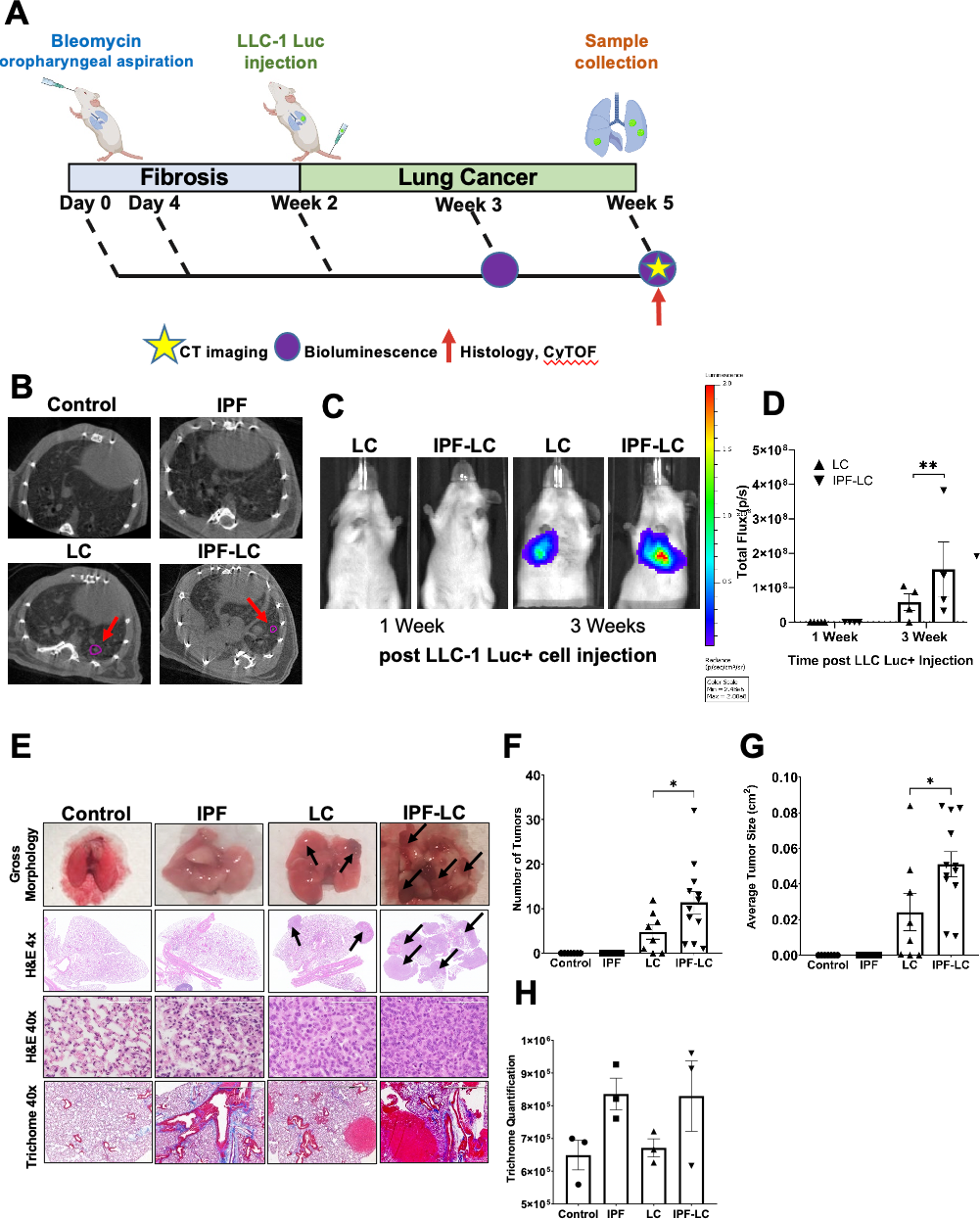
Modeling Pathogenesis of Idiopathic Pulmonary Fibrosis-Associated Lung Cancer in a Murine Model. (**A**) Schematic indicating experimental e: C57BL/6-albino mice were randomized into four experimental cohorts for all comparative studies: control, IPF, LC and IPF-LC. Mice in the control received appropriate vehicles (control), mice in the IPF group received 0.5 mg/kg and 1 mg/kg of bleomycin on day 0 and day 4, respectively, by aryngeal aspiration (OA). Mice in the LC group received LLC-1 Luc cells (1×10^6 cells/100µl) by intravenous injection into the tail vein (IV) on day 14 ice in the IPF-LC group received 0.5 mg/kg and 1 mg/kg of bleomycin (OA) on day 0 and day 4, respectively, and LLC-1 Luc cells (IV) on day 14 . oints for micro CT and bioluminescence imaging (BLI) or tissue collection are denoted by a yellow star, a purple circle and a red arrow respectively. (**B**) sentative Micro CT images from all groups at study endpoint. Tumor lesions in LC group and IPF-LC group are indicated by red arrows and circled in **C**) Representative BLI images from mice in LC and IPF-LC groups at week 1 and 3 (day 21 and day 28) post LLC-1 Luc injection. (**D**) Quantification of inescence at indicated timepoints of mice in the LC (n=4), and IPF-LC (n=4) groups. Data is presented +/− SEM and statistics were performed using one-NOVA with post-hoc Tukey (Honestly Significant Difference (HSD) test with statistical significance denoted as **p≤0.01. (**E**) Representative images of morphology, H&E and trichome staining of lung images and lung sections acquired at study endpoint at 5 weeks post first bleomycin injection. Black s indicate tumors on H&E stained lung sections of mice in LC and IPF-LC groups. (**F**) Quantification of the number of tumors in the LC and IPF-LC groups d on 1x H&E-stained lung sections presented +/− SEM. Statistical test was performed with one-way ANOVA with post-hoc Tukey HSD test and cance indicated as *p≤0.05. (**G**) Quantification of the average tumor size in the lung in LC and IPF-LC groups measured on 1x H&E-stained lung sections. s represented +/-SEM and statistical significance determined using one-way ANOVA with post-hoc Tukey HSD test and significance of *p≤0.05. (**H**) ification of collagen in trichrome staining of lung sections. No statistical difference was determined between groups.

To develop a more physiologically relevant and spontaneous model of IPF-associated lung cancer we utilized two previously described inducible transgenic models where lung cancer-associated Kras^G12D^ and fibrosis-associated TGF-alpha expression is directed to the lung through the control of reverse tetracycline transactivator by the Clara cell secretory protein promoter Scgb1a1 (CCSP) [32, 41] (**Supplemental Fig. 1**). The reverse tetracycline transactivator is expressed in club cells (rtTA), and in the presence of doxycycline, results in expression of Kras^G12D^ and/or TGF-alpha in these cells, thus inducing lung cancer or/and fibrosis. We generated four experimental cohorts: (1) control mice lacking CCSPrtTA, Kras^G12D^, and TGF-alpha, (2) LC-positive mice with CCSPrtTA and Kras^G12D^ but no TGF-alpha transgene, (3) IPF-positive mice with CCSPrtTA and TGF-alpha but no Kras^G12D^ transgene, and (4) IPF-LC mice that were positive for all three transgenes. All mice were administered doxycycline twice a week at a concentration of 0.5 mg/mL in drinking water for the duration of the 20-week experiment. As expected, expression of TGF-alpha in the lung induced fibrosis in both the IPF and IPF-LC groups and expression of Kras^G12D^ resulted in lung cancer lesions in the LC group. However, the fibrosis progressed quickly and resulted in mortality prior to the detection of lung tumors in the IPF-LC group. Statistical analysis revealed significant differences in weight changes between the IPF or IPF-LC groups when compared to control or LC groups, with IPF and IPF-LC mice exhibiting a marked loss of body weight (**Supplemental Fig. 1B**). Additionally, the IPF and IPF-LC groups exhibited significantly accelerated mortality compared to other groups, with only the LC and control groups surviving until the end of the 20-week study period (**Supplemental Fig. 1C**). As such, this spontaneous genetically engineered model of IPF-associated lung cancer, was found to be inadequate to determine the impact of fibrosis on tumor growth and aggressiveness. Future modification of this model where expression of TGF alpha and Kras^G12D^ is controlled independently of each other will be necessary to answer this question. In summary, we have developed a murine model using bleomycin and LC cells that recapitulates the pathobiology of IPF-associated lung cancer and have detected an increased tumor load in fibrotic lungs.

### Tumor progression of pulmonary lesions is associated with increased Tumor Associated Macrophages (TAMs) and decreased T cell infiltration

The tumor immune microenvironment (TIME) plays a pivotal role in tumor initiation and progression by creating an immune suppressive, tumor promoting or tumor killing microenvironment. To study the infiltrating immune cells in IPF associated lung cancer presumably contributing to tumor initiation and promotion, we utilized a previously described tissue staining method to phenotype tumor epithelial cells (CK19), macrophages (F4/80), Arginase 1 expressing immune suppressive tumor associated macrophages (TAMs), and T cells (CD3, CD8, CD4) [36]. Multiplex fluorescent immunohistochemistry was performed on murine lung sections of all four groups (control, IPF, LC and IPF-LC) (Fig. 2A). Increase in macrophage abundance or infiltration as determined by F4/80 marker expression was observed in the lungs of IPF-LC mice when compared to LC mice, indicating their contribution to accelerated tumor growth in this group (Fig. 2B). Arginase 1 positive cells, a phenotypic marker for Tumor associated macrophages (TAMs) often associated with M2 polarized immune suppressive macrophages, were detected in the IPF-LC group with a statistically significant difference to the LC group (Fig. 2C). Interestingly, a significant decrease in Cytokeratin 19 (CK19) staining of epithelial cells was observed in IPF-LC when compared to LC tumors (Fig. 2D). This may indicate a TGF dependent epithelial to mesenchymal transition that results in a reduction of cytokeratin 19 (CK19) expression, as previously described in lung epithelium [42]. When querying CD8 cytotoxic T cells specifically, no statistical difference between the LC and IPF-LC groups was observed (Fig. 2E and F), but a trend towards a decreased cytotoxic T cell abundance was observed which would support a phenotype of more rapid tumor promotion observed in the IPF-LC group and a immune suppressive tumor microenvironment. To further assess proliferation and differences in T cell abundance additional histological analyses were performed on lung sections from all experimental cohorts (control, IPF, LC and IPF-LC). Lung sections were stained for proliferation marker Ki67, but no statistical difference was observed between tumor bearing mice in the LC and IPF-LC groups (Fig. 2G and H). CD3 T cells and CD8 cytotoxic T cells were reduced in IPF-LC lung lesions and limited to the periphery of the tumor when compared to tumors in the LC group. (Fig. 2I and J). In summary, we observed an increase in macrophages with Arginase 1 expressing TAMs predominantly enriched in lungs from IPF-LC mice and a decrease in cytotoxic T cells. This finding indicates macrophage polarization to an M2 phenotype, which corroborates with our findings of increased tumor growth in lungs of IPF-LC mice.

**Figure 2:**
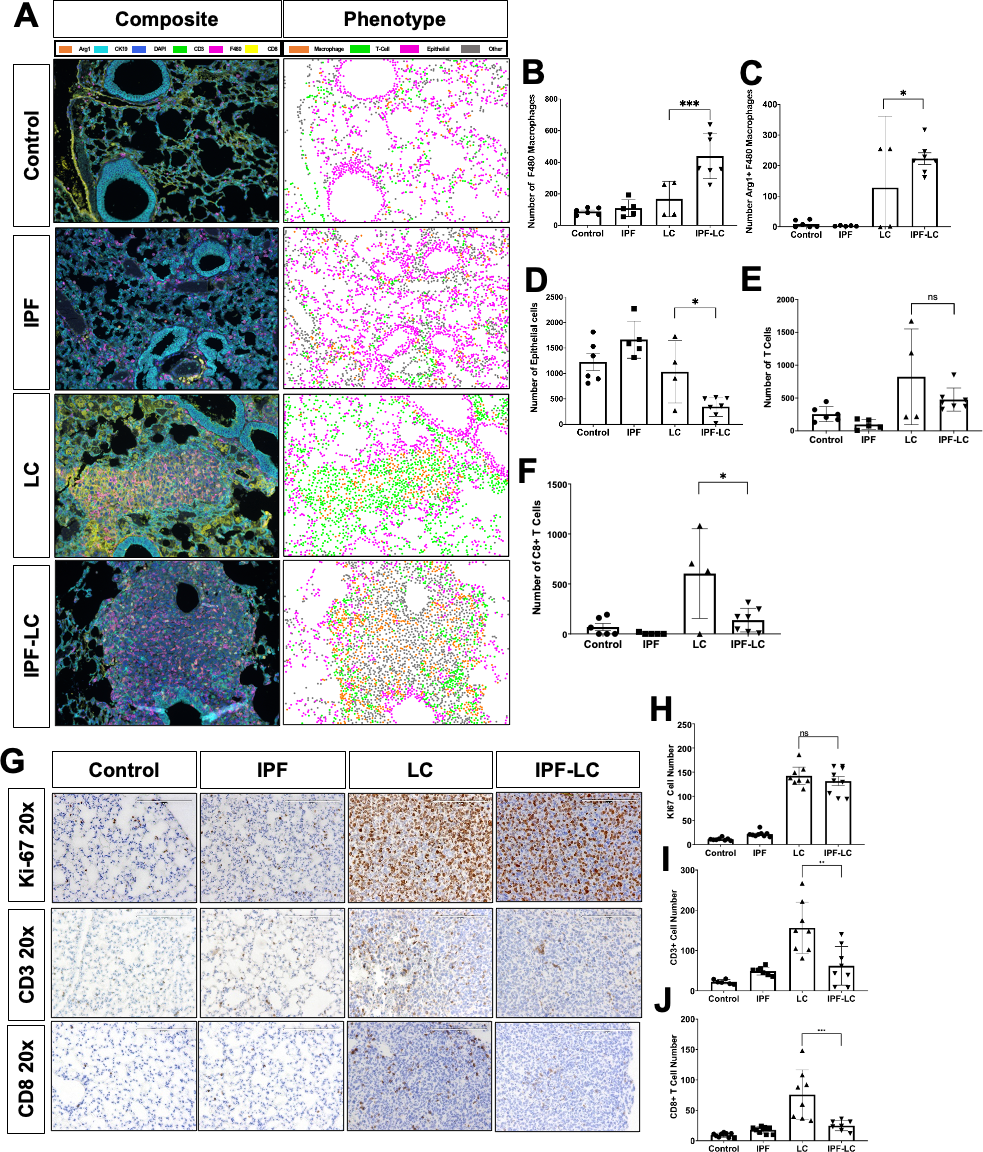
Tumor progression of pulmonary lesions is associated with increased macrophage infiltration and decrease of cytotoxic T cells. **(A)** Multiplex immunohistochemistry/immunofluorescence (mIHC/IF) performed on lung section from at least three mice per group. Representative composite images of IF (left side) are shown at 4x magnification and antibody stains are shown in corresponding colors: Arg1 (Orange), CK19 (light blue), DAPI (dark blue), CD3 (green), F4/80 (pink), CD8 (yellow). Representative images of computational phenotype are depicted on the right with cell phenotypes in corresponding color as indicated: macrophages (orange), T-cells (green), epithelial (pink), other (grey). **(B)-(F)** Quantification of /80+ macrophages, Arg1+ F4/80 macrophages, and CK19 epithelial, CD3 and CD8 positive T cells abundance of quantified composite images. Data epresented +/− SEM and statistical significance determined by one-way ANOVA with post-hoc Fisher’s Least Significant Difference (LSD) test and noted as *p<0.05, ***p<0.005 or ns for no statistical difference **(G)** Representative images (20x) of lung section in each group at endpoint stained with Ki-67, CD3 and CD8. **(H)-(J)** Quantification of Ki67, CD3+ T-cells and CD8+T-cells counted on lung section stained with corresponding antibody. Data represented +/− SEM and statistical significance determined with one-way ANOVA with post-hoc Tukey HSD test and significance of **p<0.01, *p<0.001 or ns for not statistically different.

### CD274 expression indicates reprogramming of Macrophages in lung cancer associated with IPF

To study the relative abundance of immune cells in IPF-LC, lungs from control mice, mice treated with bleomycin (IPF), mice injected with LLC-1 Luc cells (LC), or both (IPF-LC) were obtained at study end point, homogenized, and suspended as single cells for CyTOF analysis (Fig. 3A). Confirming our previous observations by mIHC, macrophages, specifically TAMs were found to be significantly elevated in the IPF-LC group when compared to lungs from IPF or LC mice (Fig. 3B and C). However, CD45+, CD3+ Ly6C high MDSCs, likely tumor associated neutrophils (TANs) showed a downward trend (Fig. 3D). No statistically significant difference in overall B, CD 4 T cell or APC/dendritic cell number was determined (see **Supplemental Fig. 2**). Interestingly, flow cytometry of dissociated lung section indicated an increase in CD274 marker expression, supporting an immune suppressive and TAM like phenotype (Fig. 3E and F). In summary, our findings suggest that reprograming of lung resident macrophages to TAMs which likely express the CD274 marker in IPF-LC group compared to the LC group supporting immune suppressive and tumor promoting niche for lung cancer progression and metastases.

**Figure 3:**
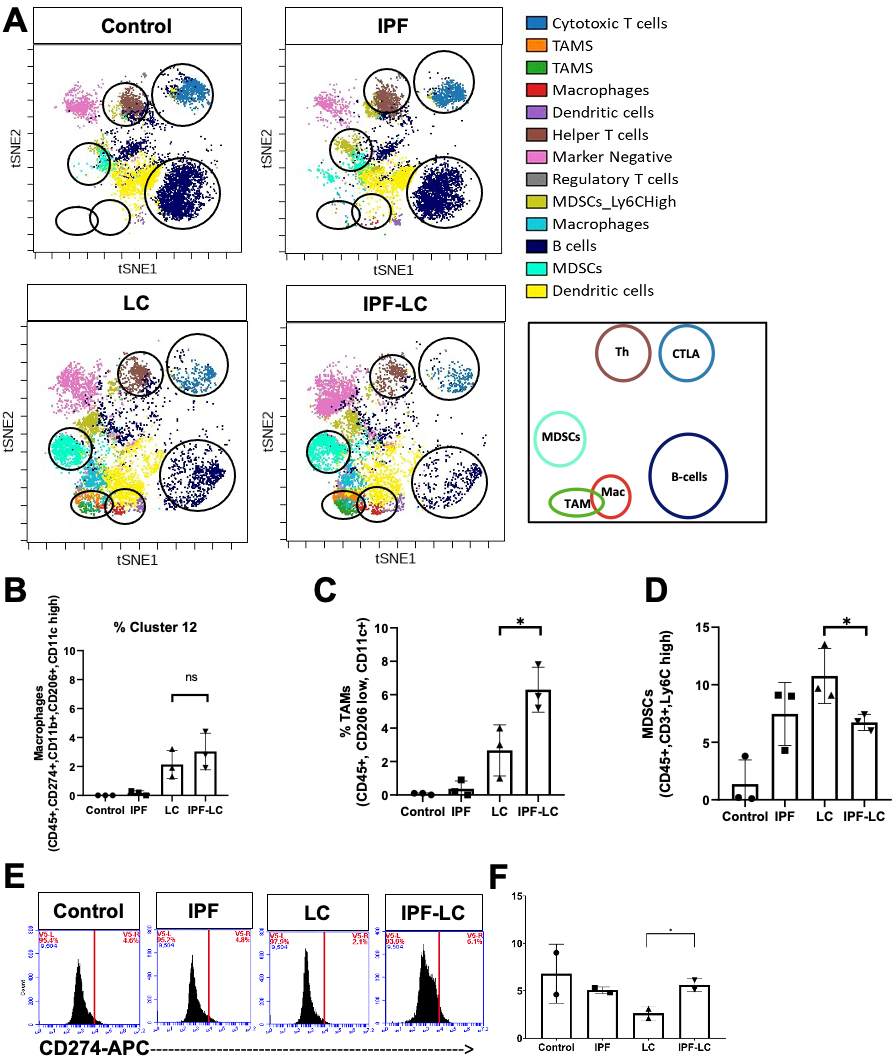
Increased Tumor associated Macrophages (TAMs) and CD274 expression in IPF-LC. **(A)** CyTOF of at 3 lungs per experimental group was performed at study endpoint (5 weeks post initial bleomycin injection). Representative tSNE plots of CyTOF analysis with specific immune cell clusters depicted and circled in corresponding colors **(B)-(D)** Quantification of abundance of cell specific expression markers for macrophages (CD45+, CD11b+, CD274+), TAMs (CD45+, CD206low, CD11c+) and MDSCs (CD45+, CD3+, Ly6Chigh), using FlowSOM-viSNE. Data +/−SEM with statistical analysis using One-way ANOVA with post-hoc Fisher’s Least Significant Difference (LSD) Test (*p.0.05, ns for no statistical difference). **(E)-(F)** Flow cytometry of lung tissue obtained from 2 mice per group at endpoint stained for CD274 marker expression using anti-CD274-APC. Statistical significance was determined using One-way ANOVA with post-hoc Fisher’s Least Significant Difference (LSD) Test (*p≤0.05).

### Identification of a unique gene signature of IPF-associated lung cancer

To gain mechanistic insight into the molecular underpinnings of IPF-LC we performed bulk RNA sequencing analysis on lung tissue obtained from control, IPF, LC and IPF-LC mice. Sequencing was performed on lung tissue from three mice per group and Principal Component Analysis (PCA) (see **Supplemental Fig. 3**) and an unbiased cluster analysis of the sequencing data was performed. A heatmap of gene log2 fold changes for the indicated comparisons shows distinct patterns between the groups when compared to control or each other (Fig. 4A). Genes with statistical significance are highlighted with intense blue or red colors based on whether they were down- or upregulated, respectively. We then compared IPF-LC to IPF and LC to expand our initial heatmap and found the log2fold change to be lower for both up- and downregulated genes when comparing these groups, as opposed to comparing IPF-LC, IPF, and LC to control (Fig. 4A). To depict overlap and shared up- and downregulated genes as well as independently regulated genes in the different groups we utilized Venn Diagrams (Fig. 4B and C). Interestingly, we identified unique gene signatures for IPF-LC and LC groups, but not for IPF, along with overlap between IPF-LC and LC groups (Fig. 4B and C **and Supplemental material**). Next, we focused on genes uniquely modulated in the IPF-LC group to better understand the molecular pathobiology of IPF-LC. The log2 fold changes for the top 25 up- and downregulated protein-coding genes in IPF-LC are shown in a heatmap comparing how the genes are expressed in LC, IPF and IPF-LC groups versus control (Fig. 4D). In summary, we have identified uniquely regulated genes in the IPF-LC groups, which may provide insight into the pathobiology of IPF-associated lung cancer.

**Figure 4:**
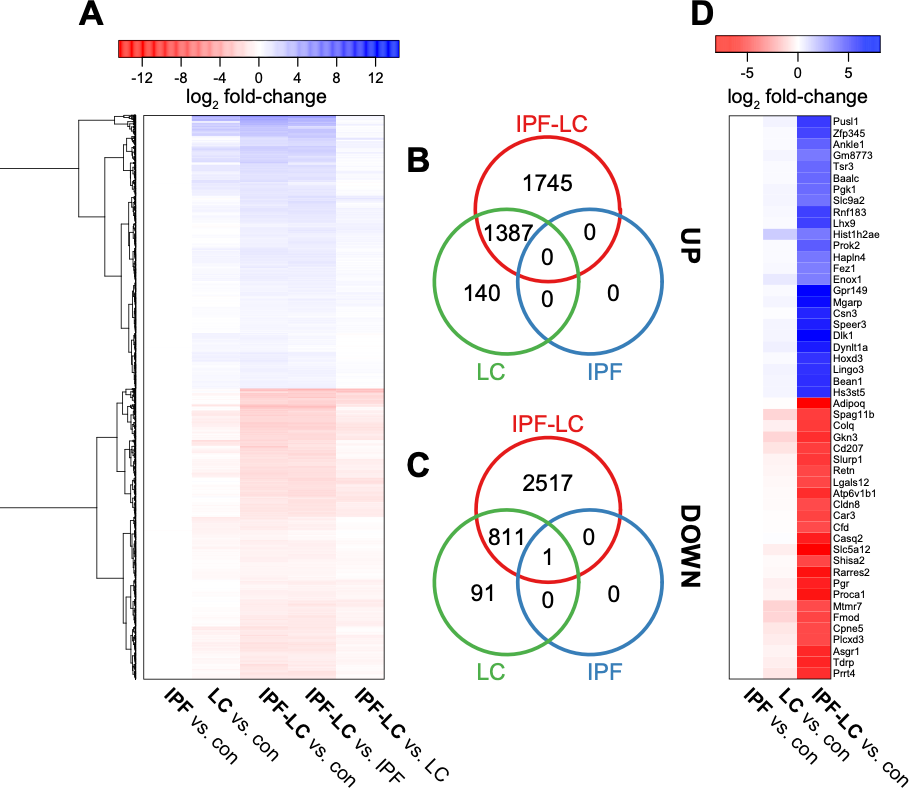
RNA Sequencing identifies genes uniquely regulated in IPF-LC. Lung tissue from three mice per group was used at endpoint for bulk RNA sequencing to evaluate differential gene expression in experimental groups. **(A)** Heatmap of gene log2 fold changes for the indicated comparisons. **(B)** and **(C)** The intersections of genes significantly differentially expressed in IPF-LC, IPF, and LC (vs control) are represented by Venn diagrams for upregulated (B) and downregulated **(C)** genes. **(D)** Heatmap of log2 fold changes for the top 25 up- and downregulated protein-coding genes in IPF-LC only. Genes with adjusted p-values < 0.05 and fold change > 1.5 were considered significantly differentially expressed.

### Pathway analysis identifies immune suppressive, secreted factors uniquely regulated in IPF-LC

To better understand the molecular pathobiology of IPF-LC and identify signaling nodes regulated in this disease, we performed GSEA Hallmark pathway analysis using all genes uniquely regulated in IPF-LC. Interestingly, but not surprisingly, we found that myc, oxidative phosphorylation, DNA repair, E2F targets and mTOR signaling pathways were upregulated and identified interferon gamma response, interferon alpha response and inflammatory response among the downregulated pathways (Fig. 5A). Querying genes in the three Hallmark pathways for interferon gamma, alpha and inflammatory responses in the four experimental groups, we found their expression to be significantly reduced in the IPF-LC group when compared to control (Fig. 5B). This was validated for a select group of genes by qRT-PCR (Fig. 5C). In summary, signaling pathway analysis and confirmatory qRT-PCR identified genes within inflammatory and interferon response pathways to be downregulated upon tumor progression in the IPF-LC group, providing evidence of an immune suppressive tumor microenvironment.

**Figure 5:**
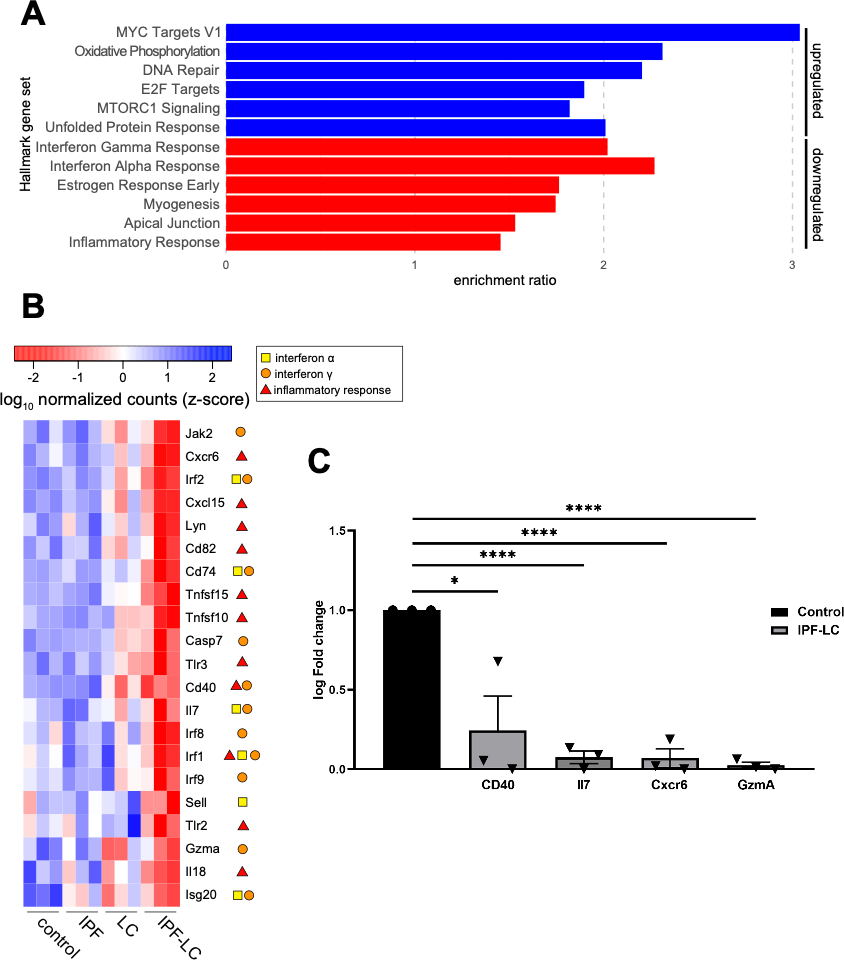
Pathway analysis identifies secreted factors uniquely regulated in IPF-LC. **(A)** Enrichment of mSigDB Hallmark gene sets (FDR < 0.05) for all IPF-LC-regulated genes (blue: upregulated; red: downregulated). **(B)** Selected genes, which appear in three indicated Hallmark gene sets depicted in **(A)** differentially expressed in IPF-LC vs control. Heatmap of log10 normalized counts (z score) for these selected genes in all groups: control, IPF, LC, IPF-LC. Symbols in legend indicate in which Hallmark gene sets they appeared (interferon gamma response, interferon alpha response, inflammatory response). **(C)** qRT-PCR of selected genes downregulated in the IPF-LC group, normalized to cyclophilin as housekeeping gene and fold change calculated compared to control. Significance was determined using unpaired student T-test of RNA obtained from lung tissue of three mice per control and IPF-LC group. * Denotes statistical significance of p<0.05, *** and p<0.005.

### Loss of Cytokeratin 7 marks Epithelial Mesenchymal Transition (EMT) in IPF-LC

EMT, the process by which epithelial cells lose their epithelial abilities and become mesenchymal cells and increase their expression of cancer stemness genes, is observed in many malignancies, including lung cancer, when tumors progress. We queried known EMT gene expression to identify a potential epithelial to mesenchymal transition in the IPF-LC group denoting more aggressive tumor growth. Genes that are typically expressed in epithelial cell compartments, like Cytokeratin 7/Krt7, Mucin 1/Muc1 and E Cadherin/Cdh1, were downregulated in the IPF-LC group, whereas genes that denote a more mesenchymal phenotype, like Vimentin/Vim, N-Cadherin/Cdh2 and Fibroblast specific protein (FSP)-1 (also called S100A4), were upregulated (Fig. 6A). Next, we performed IHC on lung tissue obtained from the four different experimental groups and found that Cytokeratin 7 expression (CK7) was completely lost in the IPF-LC group (Fig. 6B), indicating a mesenchymal, more aggressive, and potentially invasive and metastatic phenotype. In summary, the progression of lung cancer observed in IPF-associated lung cancer may in part be due to EMT.

**Figure 6:**
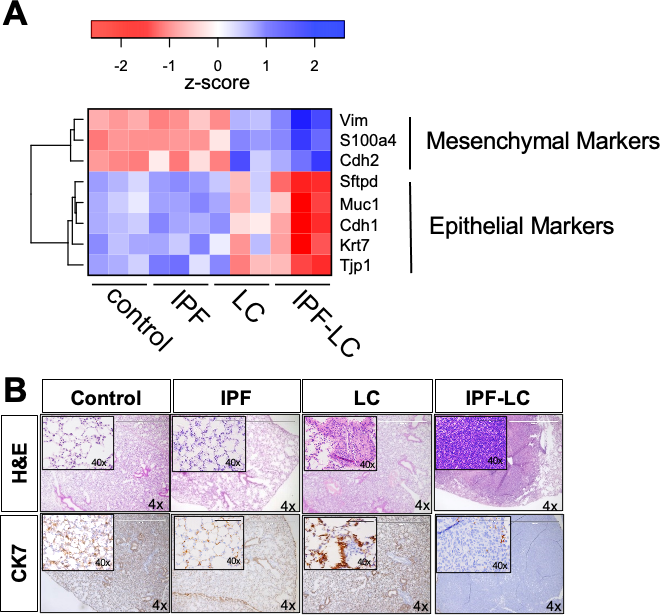
Pronounced EMT in IPF associated lung cancer. RNA sequencing data as described in Figure 4 was used to query genes important in EMT. **(A)** Heatmap of log10 normalized counts (z score) for selected mesenchymal and epithelial marker genes in all groups: control, IPF, LC, IPF-LC. **(B)** Representative images (4x, (40x inset)) of lung section in each group stained with H&E (top) and Cytokeratin 7 (bottom).

## Discussion

In this study we developed and explored a new mouse model for idiopathic pulmonary fibrosis (IPF)-associated lung cancer to gain mechanistic insight into the link between IPF and lung cancer and to identify potential therapeutic targets and develop future treatment options for patients with IPF-associated lung cancer (IPF-LC). Our model combined the commonly used bleomycin model for IPF with the Lewis lung adenocarcinoma (LLC-1 Luc) model to improve understanding of the lung microenvironment in the development of IPF-LC. We identified an increase in immune cell infiltration that likely contributes to a tumor promoting and immune suppressive microenvironment in IPF-LC but not in lung cancer or IPF alone. Our findings indicate that macrophages, particularly tumor associated Arginase + macrophages, may communicate with T cells to produce a tumor promoting environment that results in growth and proliferation as assessed by pre-clinical CT and BLI and histological assessment. Furthermore, we identified a unique gene signature in IPF-LC that provides biological insight into the pathobiology of this disease and may provide future biomarkers and therapeutic targets.

Development of a new mouse model for IPF-LC provides an important tool to interrogate the role of IPF and other pre-existing lung diseases in the development of lung cancer. While our findings utilizing this model contribute to the understanding of IPF-LC and provide an opportunity for future mechanistic work and testing of therapeutic interventions, this model has some limitations. In bleomycin-induced pulmonary fibrosis models, the rapid onset of disease and apparent resolution within a few weeks is not a faithful representation of human pulmonary fibrosis, which develops over many years and does not resolve on its own [43]. Moreover, our model specifically recapitulates only some aspects of idiopathic pulmonary fibrosis; for simplicity we have used ‘IPF’ as nomenclature. However, while our model does not fully recapitulate the human disease, it exhibits key molecular characteristics of human IPF, such as intra-alveolar buds, mural incorporation of collagen, and obliteration of the alveolar space [44]. The combination of the bleomycin model with the LLC-1 lung tumor metastasis model, which bears the Kras^G12C^ mutation, has previously been explored [45]; however, this has not provided insight into the composition of the tumor microenvironmental or transcriptomic changes in IPF-associated lung cancer. The LLC-1 model has its limitations as well, as it only recapitulates lung cancer with the Kras^G12C^ mutation, but not cancer with other Kras or co-occurring mutations in other oncogenes or tumor suppressors that may affect the development of an immune suppressive, tumor promoting microenvironment differently [30, 46]. At the same time, it is important to note that Kras^G12C^ mutant lung adenocarcinoma is one of the most aggressive forms of lung cancer, with a mortality of 60% [47], and although it likely doesn’t represent all IPF-associated lung cancers, it is representative of the majority of lung cancers. Furthermore, other mutant Kras cell lines can be developed to generate a similar IPF-LC model with other oncogenic mutations. Furthermore, our study explored the use of a genetically engineered mouse model to recapitulate lung cancer associated with pulmonary fibrosis. In this model, KrasG12D and TGF-alpha were expressed in club cells of the lung to simultaneously promote the development of fibrosis and lung cancer. However, the fibrosis progressed quickly and resulted in mortality prior to the detection of lung tumors in the IPF-LC group. Herzog et al. recently developed a bleomycin-induced lung fibrosis model in combination with a genetically engineered mouse model of NSCLC and showed, similarly to us, that fibrosis exacerbated lung cancer progression [25]. Notably, progression was halted when TGF-beta signaling was inhibited [25].

In summary, despite the limitations described for our IPF-LC model, this model provides an opportunity to interrogate and therapeutically modulate both the TME and signaling pathways regulated in IPF-associated lung cancer and demonstrates that pre-existing lung disease contributes to an elevated tumor progression.

Our study findings utilizing multiplex immunohistochemistry identified that F4/80 positive macrophages were more abundant in lungs of mice with IPF-LC and just lung cancer (LC). Importantly, these F4/80 macrophages in the IPF-LC group showed a significant increase in Arginase 1 expression, which is a marker of tumor associated macrophages (TAMs), also referred to as M2-like macrophages and immunosuppressive and tumorigenic functions [48]. TAMs affect patient response to chemo-, immuno- and radiotherapies [49] and TAM abundance correlates with aggressiveness and metastatic potential of tumors, which has been demonstrated in pre-clinical mouse models and clinical studies [50]. Specifically, TAMs play a tumor promoting role because of their secretion of growth factors, like vascular endothelial growth factor (VEGF), that facilitate tumor cell motility and extracellular matrix remodeling. Furthermore, TAMs express immunosuppressive factors, like PDL1, that prevent immune cells from recognizing tumor cells [49]. Interestingly, CD8 T cells appeared to be downregulated in the IPF-LC group, supporting a more immune suppressive and tumor promoting microenvironment. It remains to be investigated whether the IPF induced lung microenvironment results in a polarization of lung resident alveolar macrophages repolarized by disseminated tumors cells that seed in the lung, or in a recruitment of distant, bone marrow derived macrophages that infiltrate the lung to provide a pre-metastatic niche. Moreover, TAMs provide viable targets for immune therapies [51]. Targeted therapies to inhibit or reprogram TAMs, that may be used in combination with other therapies, are being developed to increase prognosis in cancer patients [52]. In summary, our data indicates that pre-existing lung disease, like fibrosis, increases the infiltration or polarization of macrophages, specifically M2 TAMs that express Arginase 1. As such, these macrophages contribute to the high mortality rates and the low 5-year survival observed in patients with IPF-LC compared to lung cancer patients.

Our CyTOF analysis allowed us to gain a better understanding of the composition of immune cells in the tumor microenvironment in all four experimental groups. Interestingly, the abundance of B cells decreased in the lung cancer group and in the IPF-LC group when compared to controls or IPF. B cells are understudied in the tumor microenvironment but are also thought to have immune suppressive functions [53]. Although we did not observe a major difference in B cell abundance between LC and IPF-LC it remains to be investigated whether their regulatory function is differentially modulated. We found that macrophages and TAMs, but not Ly6C myeloid-derived suppressor cells (MDSC), were more abundant in the IPF-LC group, further supporting an immune suppressive tumor promoting microenvironment in this group when compared to LC alone. It is well established that MDSC exert their functional role of immune suppression through multiple mechanisms that include depletion of amino acids such as cystine and arginine [54, 55], production of nitric oxide (NO) and reactive oxygen species (ROS) [56], increased production of interleukin (IL)-10 and transforming growth factor (TGF)-β [55, 57], secretion of angiogenic factors [58, 59], production of cytokines, growth factors, matrix proteases and upregulation of programmed death-ligand 1 (PD-L1) [60]. Using flow cytometry, we observed an overall upregulation of PD-L1 expression in the IPF-LC group, but this may be due to expression on other myeloid derived immune cells. Because MDSC are known to regulate a plethora of immune modulatory processes, including T cell migration, T cell death, tumor neovascularization and tumor growth, we were surprised to notice a downward trend of MDSCs in the IPF-LC group. Thus, future studies will include a detailed characterization of tumors associated with IPF to evaluate the role of MDSCs, or tumor-associated neutrophils (TANs), in tumor progression in this group. Furthermore, as discussed above, the identification of TAMs and MDSCs as major players in modulating an immunosuppressive, tumor promoting microenvironment in IPF-LC may provide future opportunities for therapeutic intervention as it is being explored in other malignancies [61].

In addition, we observed trends of downregulation of T cells and dendritic cells in the IPF-LC group compared to LC alone. Specifically, we observed a downregulation of CD8+ cytotoxic T cells and CD4+ helper T cells in addition to the downregulation of antigen presenting dendritic cells. This further supports a more immunosuppressive TME in IPF-LC tumors when compared to LC as both lymphocytes are required for a cell-mediated immune response in malignancies [62].

In fact, a significant decrease in CD4+ T helper cells was observed in IPF-LC. This is interesting, as CD4 T helper cells play an important role in maintaining effective anti-tumor immunity [63] and can differentiate into effector cells needed in the anti-tumor response. Additionally, CD4+ T helper cells primarily mediate anti-tumor immunity by assisting CD8+ cytotoxic T cells and antibody responses by providing stimulus for priming CD8+ T-cells [64]. Similarly, the anti-tumorigenic role of CD8+ cytotoxic T-cells, their interaction with CD4 T cells and other immune cells and their association with better patient outcome has been well established and our findings of a reduced CD8 and CD4 T cell abundance in IPF-LC was therefore not surprising [65–71]. In summary, the noted downregulation of CD4, CD8 and dendritic cells in the IPF-LC group when compared to the lung cancer group further supports our finding of a tumor promoting and immunosuppressive TME that leads to a worse prognosis and decrease in the 5-year survival rate in patients with IPF-associated lung cancer.

In addition to identifying key immune cells that may be contributors to an immune suppressive and thereby tumor proliferative microenvironment in IPF-LC, we identified genes and signaling pathways that are uniquely modulated in this disease, providing further insight into its pathogenesis. We identified overlap in gene regulation between LC and IPF-LC, but not between LC and IPF or IPF-LC and IPF. Our expectation was to find significant overlap between the LC and IPF-LC groups given the significant tumor cell seeding and tumor cell proliferation in both; however, we were surprised that we were unable to identify genes uniquely regulated in the IPF group. The relatively low dose of bleomycin and fast resolution of the fibrosis phenotype in the bleomycin-induced model may be an explanation for these findings. Nevertheless, identification of this unique gene signature detected in IPF-LC provides insight into the signaling nodes regulated in IPF-associated lung cancer and may provide future therapeutic targets. In fact, the downregulation of the inflammatory response and interferon gamma and alpha mediated signaling of CD40 is interesting, as downregulation of CD40 has been shown to contribute to accumulation of MDSC in other cancers [72]. Furthermore, downregulation of granzyme A may indicate lack of cytotoxic T cells, which typically produce granzymes to kill aberrant cells, thereby promoting tumor growth [73]. This finding corroborates with our data indicating reduction in CD8 T cells and increased abundance of TAMs.

Lastly, our transcriptomic analysis of the four experimental groups demonstrated that IPF-associated lung cancer is accompanied by an epithelial to mesenchymal transition, thereby likely contributing to a more aggressive tumor promoting and immune suppressive microenvironment. Interestingly, bleomycin-induced fibrosis in lungs is often associated with increased Wnt 1 signaling, which initiates EMT [74]. During EMT, epithelial cells are reprogrammed to express Vimentin instead of Cytokeratin, the characteristic intermediate filaments in epithelial cells, which was lost in IPF-associated lung cancer group when compared to lung cancer alone. The gradual loss of E-Cadherin, which typically is accompanied by simultaneous gain of N-Cadherin, is considered a hallmark of EMT and commonly referred to as the “Cadherin switch” [75] and was observed in the IPF-LC group. Our findings of a more mesenchymal phenotype in the IPF-LC group were further corroborated by histological findings indicating a remarkable reduction in cytokeratin 7 staining. These findings may indicate that lung disease, especially fibrosis, caused by environmental causes, infections, or genetic predisposition, or induced by treatment, may trigger EMT in pre-cancerous cells and/or provide a niche for circulating cancer cells. As bleomycin is utilized in the treatment of several types of malignancies, including certain types of lymphoma, the long-term adverse effects should be considered [76].

In summary, we have developed a novel mouse model that recapitulates aspects of bleomycin-induced fibrosis-associated lung cancer and identified an immune suppressive TME characterized by an increased infiltration of macrophages, TAMs and MDSCs, but a decrease in CD4+ T-helper cells/CD8+ cytotoxic T-cells in lungs of IPF-LC mice, likely contributing to more aggressive tumor growth. Furthermore, we identified significant downregulation of pathways associated with responses to interferon gamma and alpha and inflammation, indicating that immune suppression may be widespread in lungs with associated fibrosis, thereby promoting tumor metastases, invasion and proliferation. In summary, our study findings provide rationale for targeting disease associated immune cells and signaling pathways to develop future therapeutic regimens for IPF-associated lung cancer.

## Acknowledgments

The authors wish to thank Michelle Paulsen of the Ljungman Lab for assisting with Bru Sequencing.

## Data sharing

The datasets supporting the current study are available from the corresponding author upon request and available in the NIH Gene Expression Omnibus database (GEO). RNA raw data files and analysis was deposited with GEO accession number GSE224134. Material or data that require a Material Transfer Agreement (MTA) can be provided by the University of Michigan pending scientific review and the execution of an MTA negotiated by the university’s Office of Technology Transfer. Requests for data that require an MTA should be submitted to the corresponding author, Dr. Stefanie Galban.

