## Supplementary Figures for "Modeling Molecular Pathogenesis of Idiopathic Pulmonary Fibrosis-Associated Lung Cancer in Mice"

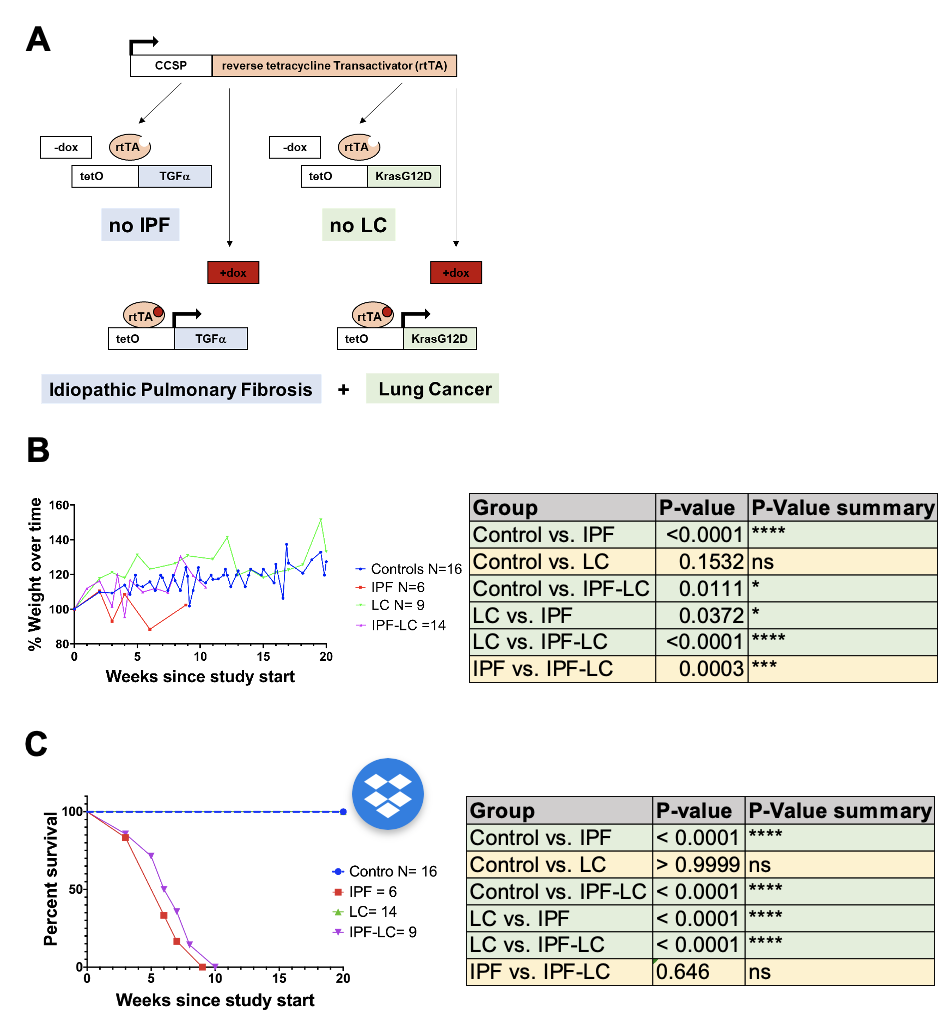

**Supplemental Figure 1: Transgenic Mouse IPF-LC model development.** (**A**) Schematic description of the transgenic mouse model used in the study. Expression of reverse tetracycline transactivator (rtTA) is controlled by the Clara cell secretory protein (CCSP) promotor in Club cells (formerly Clara cells). Addition of Doxycycline to the drinking water of mice results in activation of TGF alpha and oncogenic Kras (KrasG12D), which are under the control of Tetracycline Operon (TetO). (**B**) Weight change and (**C**) Kaplan-Meier Survival analysis of mice in control, IPF, LC and IPF-LC groups. Weights were analyzed with ordinary one-way ANOVA with Tukey's multiple comparison test and survival curves were analyzed using the Log Rank (Mantel-Cox) test. No statistical difference was determined between IPF and IPF-LC groups.

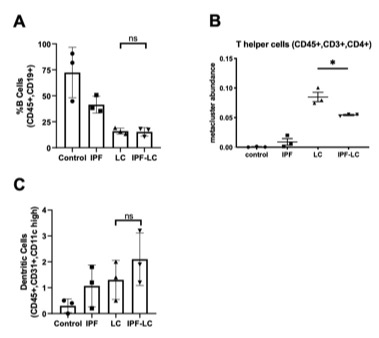

**Supplemental Figure 2: Expression of immune cell surface markers by CyTOF.** CyTOF of at least 3 lungs per experimental group was performed at study endpoint (5 weeks post initial bleomycin injection)**. (A-C)** Quantification of abundance of cell specific expression markers for B cells (CD45+, CD19+), T helper cells (CD45+, CD3+, CD4+), and dendritic cells (CD45+, CD31+, CD11c high) using FlowSOM-viSNE. Data +/-SEM with statistical analysis using One Way ANOVA with post-hoc Fisher's Least Significant Difference (LSD), n.s. denotes no significance.

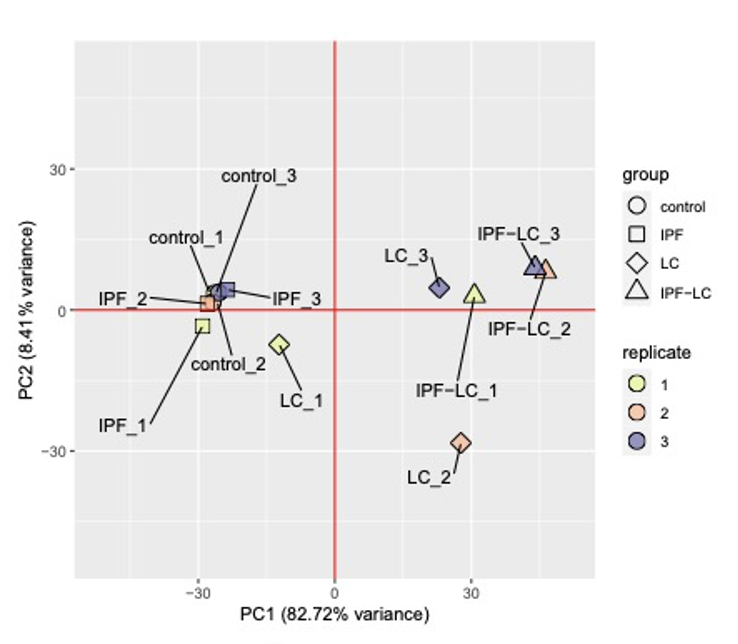

**Supplemental Figure 3:** PCA biplot (PC1 and PC2) of VST-normalized gene counts for the top 500 most variable genes. Individual samples are labeled. Groups are represented by shape (control: circle; IPF: square; LC: diamond; IPF-LC: triangle). Replicates are represented by color (1: yellow; 2: orange; 3: blue).
