## Supplementary Tables for "Modeling Molecular Pathogenesis of Idiopathic Pulmonary Fibrosis-Associated Lung Cancer in Mice"

**Supplemental Table 1:** stains and antibodies used for mIHC.

| **#** | **Antigen/Target** | **Company** | **Catalog #** |
| --- | --- | --- | --- |
| 1 | DAPI | Akoya Bioscience | FP1490 |
| 2 | CK19 | Max Plank Institution | Troma III |
| 3 | CD3 | Dako | A0452 |
| 4 | CD8 | Cell Signaling | 98941 |
| 5 | F480 | abcam | Ab6640 |
| 6 | Arg1 | Cell Signaling | 93668 |

**Supplemental Table 2**: antibody panel used for CyTOF analysis.

| **#** | **Antibody Target (Antigen)** | **Catalog # (Fluidigm)** | **#** | **Antibody Target (Antigen)** | **Catalog # (Fluidigm)** |
| --- | --- | --- | --- | --- | --- |
| 1 | Ly-6C | 3150010B | 10 | CD140a | 3148018B |
| 2 | Ly-6G | 3141008B | 11 | CD3e | 3152004B |
| 3 | CD11b (Mac-1) | 3143015B | 12 | TCRgd | 3159012B |
| 4 | CD45 | 3089005B | 13 | CD152 (CTLA-4) | 3154008B |
| 5 | CD8a | 3168003B | 14 | CD11c | 3209005B |
| 6 | CD206 (MMR) | 3169021B | 15 | CD161 (NK1.1) | 3170002B |
| 7 | CD19 | 3149002B | 16 | CD274 (PD-L1) | 3153016B |
| 8 | CD4 | 3145002B | 17 | EPCAM (*NEW) | 3165013B |
| 9 | F4/80 | 3146008B | 18 | CD31 (PECAM-1) | 3165013B |
